# Capturing single-molecule properties does not ensure accurate prediction of biomolecular phase diagrams

**DOI:** 10.1101/2024.10.24.619983

**Authors:** Alejandro Feito, Ignacio Sanchez-Burgos, Antonio Rey, Rosana Collepardo-Guevara, Jorge R. Espinosa, Andres R. Tejedor

## Abstract

Intracellular liquid-liquid phase separation of proteins and nucleic acids represents a fundamental mechanism by which cells organise their components into biomolecular condensates that perform multiple biological tasks. Computer simulations provide powerful tools to investigate biomolecular phase separation, offering microscopic insights into the physicochemical principles that regulate these systems. In this study, we investigate the phase behaviour of the low-complexity domain (LCD) of hnRNPA1 and several mutants via Molecular Dynamics simulations. We systematically compare the performance of five state-of-the-art residue-resolution coarse-grained protein models: HPS, HPS-cation-*π*, CALVADOS2, Mpipi, and Mpipi-Recharged. Our evaluation focuses on how well these models reproduce experimental coexistence densities and single-protein radii of gyration for the LCD-hnRNPA1 set of mutants. While most models yield similar intramolecular contact maps and reasonable estimates of the single-protein radius of gyration compared to in vitro measurements, only Mpipi-Recharged, Mpipi, and CALVADOS2 accurately predict phase diagrams that align with experimental data. This suggests that force field parameterizations optimized solely to reproduce single-protein properties may not always capture the phase behaviour of protein solutions. Additionally, our findings reveal that some residue-resolution coarse-grained models can lead to significant discrepancies in predicting the roles of individual intermolecular interactions, even for relatively simple intrinsically disordered proteins like the low-complexity domain of hnRNPA1. Our work highlights the importance of balancing both single-molecule and collective properties of proteins to accurately predict condensate formation and material properties.

## I. INTRODUCTION

Liquid-liquid phase separation (LLPS) is a crucial mechanism exploited by cells to organise their matter in time and space, enabling them to carry out numerous biological functions^1–4^. LLPS of biomolecules, such as multivalent proteins and nucleic acids^5–7^ enable the formation of membraneless organelles ranging from P granules^2,8^ or ribonucleoprotein granules^9,10^, located at the cytosol, to paraspeckles^11–13^ or Cajal bodies^14,15^ which are present in the cell nucleus ^16,17^. Crucially, biomolecular condensates are central to regulate diverse cellular functions such as genome silencing^18,19^, cell signaling^20–22^, compartmentalisation^23–25^, or buffering protein concentration^23,24^ among many others^26,27^.

Proteins that are either intrinsically disordered—e.g. DEAD-Box helicase 4 (DDX4) protein^28,29^, R12^30^—or multi-domain proteins that contain both globular regions and low complexity domains (LCDs)—e.g. *fused in sarcoma* (FUS)^31–33^, the TAR DNA-binding protein of 43 kDa (TDP-43)^33–35^, the heterogeneous nuclear ribonucleoprotein A1 (hnRNPA1)^36,37^, or heterochromatin protein 1 (HP1)^38–40^—represent principal constituents of biomolecular condensates formed via LLPS^41–43^. The flexibility of the disordered regions allows proteins to efficiently adapt their conformation inside condensates and establish multiple intermolecular contacts that increase their thermodynamic stability^8,43–46^. As a consequence of the high density of connections they form, some LCDs can undergo disorder-to-order structural transitions^47,48^ inside biomolecular condensates. These structural transitions have been directly related to progressive stabilization and rigidification of biomolecular condensates against dissolution, giving rise to aberrant solid toxic assemblies which are connected to neurodegenerative disorders such as amyotrophic lateral sclerosis (ALS)^49–51^, frontotemporal dementia (FTD)^10,51,52^, or Parkinson^53–55^ among others^17,56^. Hence, unravelling the intricacies of biomolecular condensate formation and the variation over time of the intermolecular liquid network that sustains them can expand our microscopic understanding of such diseases^33,57–59^.

Extensive experimental studies have been devoted to understanding the fundamental thermodynamic and physicochemical factors underlying the transition from homogeneous protein solutions to phase-separated condensates^16,37,45,57,60,61^. In vitro studies have highlighted the critical role of the amino acid composition and patterning on the protein’s propensity to undergo phase separation^62–66^. These studies emphasize how a few key mutations can dramatically alter the material properties of condensates^62,64^. Notably, these studies have revealed that aromatic residues such as phenylalanine, tryptophan, and tyrosine play a pivotal role in driving phase separation in prion-like domain proteins by forming *π* −*π* and cation-*π* interactions^28,31,63,67,68^. Furthermore, electrostatic interactions have been shown to significantly regulate condensate formation, influenced by salt concentration, ion identity^69^, or the presence of highly charged biomolecules such as RNA^5^, DNA^18,70^, or highly charged poly-peptide repeats^71,72^. However, the exact mechanisms by which dominant network connectivity and interactions within condensates vary with solution conditions, condensate composition, amino acid sequence patterning, and protein structure cannot be uncovered solely by experiments^59,73^.

Computer simulations represent a fundamental tool to gain microscopic information on biomolecular phase separation^59,74^. Different simulation approaches have been adopted to tackle this problem, including atomistic simulations^69,75–77^, lattice-based simulations^78–80^, sequence-dependent residue-resolution coarse-grained models^28,74,76,81,82^ and minimal models^83–88^. Among the many different approaches, sequence-dependent coarsegrained models have emerged as the most efficient computational method which still captures the chemical identity of the different residues^74,89^. Residue-resolution coarse-grained models have been extensively used to unravel the underlying factors leading to biomolecular phase separation, paying particular attention to the effect of protein length^90–92^, protein conformation^81,93–95^, patterning of strong versus weakly interacting residues^28,68,76,89,96^, and the role of nucleic acids^87,88,90,92,97^. Importantly, residue-resolution models can provide molecular and thermodynamic information underlying biomolecular phase separation that is hard to measure experimentally^59,74^, such as potential energy landscapes, protein structural conformational ensembles, or relative frequency of amino acid pair interactions^74,81,90,92,98^. In that respect, the availability of residue-resolution coarse-grained models that can describe the phase behaviour of proteins and nucleic acids with near-quantitative accuracy is highly desirable since it can significantly contribute to advancing our understanding of the multiple factors dictating biomolecular phase transitions^59^.

In this work, we explore the ability of five different state-of-the-art residue-resolution coarse-grained models to capture the phase behaviour of the LCD of hnRNPA1—a multifunctional protein involved in RNA binding and splicing^99,100^, transcriptional regulation^100,101^, and stress response^102,103^—and five mutants of this protein with different types of aromatic and charged residues. Specifically, we employ the HPS^89^, HPS-cation-*π*^28^, CALVADOS2^81^, Mpipi^104^ and Mpipi-Recharged^105^ models (details of the corresponding interactions are included in Section SI of the Supplementary Material (SM), while simulation details are described in Section SII). We test the predictions of each model by computing temperature–concentration phase diagrams and comparing with the experimental data^63,65^ to benchmark their predictive capacity. We do so for the six studied proteins, and rationalise the outcome based on the differences in sequence composition. Subsequent analysis is focused on the effect of the amino acid contacts, both at intra and intermolecular level, as well as on the single-protein radius of gyration (R_*g*_) and the *θ* temperature. Strikingly, we find that while the intramolecular contact maps and R_*g*_ values predicted by all models are alike, and reasonably consistent with the experimental values^63^, the phase diagrams and intermolecular contact maps enabling phase-separation for hnRNPA1-LCD notably differ from each other depending on the model. Hence, single-protein properties such as the radius of gyration may not always be sufficient as the sole metric for benchmarking model performance or assessing the predictive capability of force fields. Overall, our results emphasize the critical need to strike the right balance between *π*–*π*, cation–*π*, and electrostatic interactions for accurately modelling condensate phase behaviour—an ongoing challenge in biomolecular simulations.

## II. PREDICTED COEXISTENCE LINES OF HNRNPA1-LCD AROMATIC-RICH MUTANTS

The stability and material properties of biomolecular condenstates depend strongly on multiple parameters such as temperature^29,63^, pH^106^, salt concentration^29,30,69,75^, or protein structural sequence and composition ^63,64^. Among these factors, amino acid composition plays a central role not only in influencing phase behaviour but also in shaping protein folding and structure. These, in turn, determine how proteins interact with their environment and, ultimately, govern their biological function^66,107–109^. Here, we compare the capacity of different coarse-grained models in capturing the impact of aromatic and charge residue mutations. For this, we compute phase diagrams of the wild-type (WT) hnRNPA1-LCD protein and five mutants—allF, allY, -9F+3Y, -3R+3K and -6R+6K. hnRNPA1-LCD (henceforth referred as WT+NLS since it also contains a nuclear localization signal; NLS) is an intrinsically disordered protein (IDP) region of 135 residues which contains numerous charged (12 positively charged residues with 10 arginines and 2 lysines; and 4 negatively charged glutamic acid residues) and aromatic (8Y and 12F) residues (Fig. 1A). The mutants studied here alter the balance between aromatic and charged amino acids or change the types of aromatic residues. For instance, one variant replaces all tyrosines (Y) with phenylalanine (F) (reaching to 19F, Fig. 1B and Table S2 for the allF sequence), another changes all F residues to Y (reaching to 19Y, Fig. 1C and Table S2 for the allY sequence), and a third one increases the abundance of tyrosine with respect of phenylalanine (−9F+3Y: reaching 10Y and 3F, Fig. 2A). The remaining variants (see Table S2) modify the presence of arginines and lysines in the sequence (reaching 7R and 5K (Fig. 2B) for -3R+3K, and 4R and 8K (Fig. 2C)) for the -6R+6K sequence.

**FIG 1.**
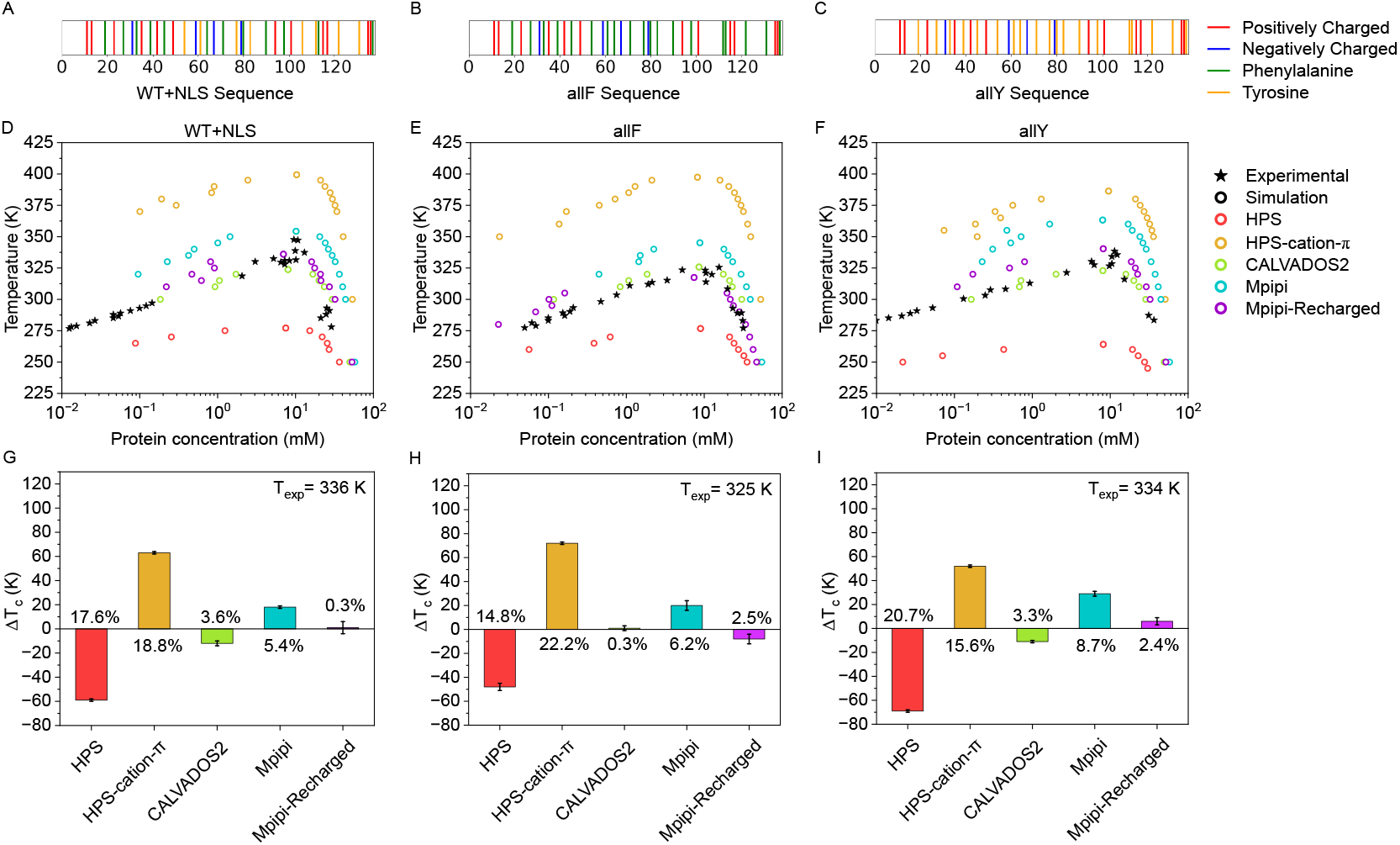
Schematic representation of the amino acid composition in the WT+NLS (A), allF (B), and allY (C) sequences. Positively charged amino acids are depicted by red lines, negatively charged amino acids in blue, phenylalanines in green, and tyrosines in orange. Experimental phase diagrams ^65^ represented by solid stars *vs*. simulated phase diagrams represented by empty circles using different models HPS, HPS-cation-*π*, CALVADOS2, Mpipi, and Mpipi-Recharged as indicated in the legend for hnRNPA1-LCD variants: WT+NLS (D), allF (E), and allY (F).Difference between simulated and experimental critical temperatures (Δ*T*_*c*_) for the different studied models represented by the same colors as in the upper panels for the sequences WT+NLS (G), allF (H), and allY (I). The relative error in percentage compared to the experimental temperature is described on each bar. Critical temperatures obtained from simulations and experimental values taken from Ref.^63,65^ can be found in Table S3 of the SM.

**FIG 2.**
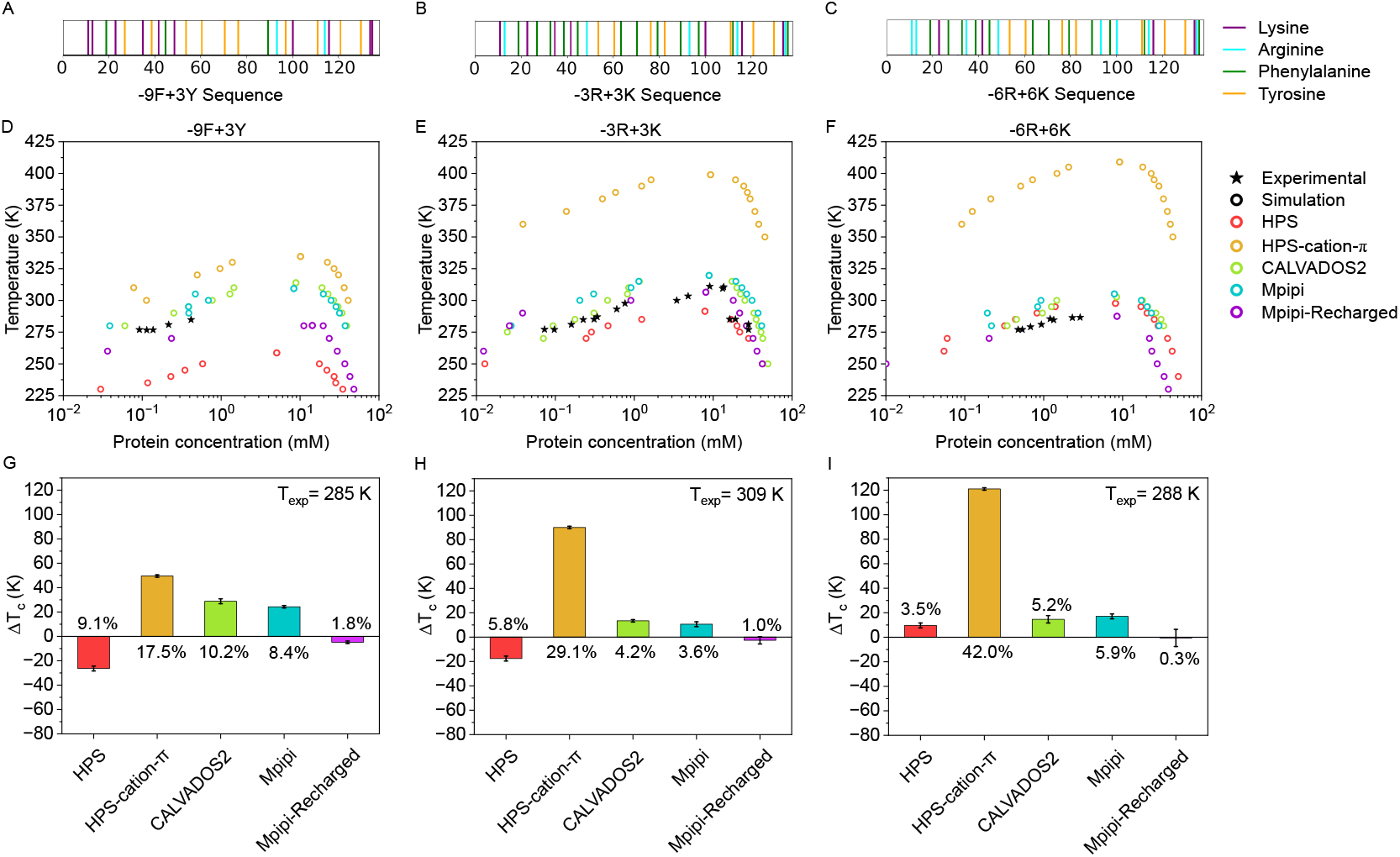
Schematic representation of the amino acid composition in the -9F+3Y (A), -3R+3K (B), and -6R+6K (C) sequences. Lysines are depicted by purple lines, arginines in cyan, phenylalanines in green, and tyrosines in orange. Experimental phase diagrams ^63^ represented by solid stars *vs*. simulated phase diagrams represented by empty circles using different models HPS, HPS-cation-*π*, CALVADOS2, Mpipi, and Mpipi-Recharged as indicated in the legend for the hnRNPA1-LCD variants: -9F+3Y (D), -3R+3K (E), and -6R+6K (F). Difference between simulated and experimental critical temperatures (Δ*T*_*c*_) for the different studied models represented by the same colors as in the upper panels for the sequences -9F+3Y (G), -3R+3K (H), and -6R+6K (I). The relative error in percentage compared to the experimental temperature is described on each bar. Critical temperatures obtained from simulations and experimental values taken from Ref.^63,65^ can be found in Table S3 of the SM.

Alshareedah et al.^65^ recently studied the phase behavior of WT+NLS and its aromatic variants, examining how these mutations influence the viscoelastic properties of condensates. This work builds on earlier studies that focused on the effects of these mutations on phase boundaries^63^. In this study, we evaluate the temperature-protein concentration phase diagrams of six sequences: WT+NLS (Fig. 1D), allF (Fig. 1E), allY (Fig. 1F), -9F+3Y (Fig. 2D), -3R+3K (Fig. 2E), and -6R+6K (Fig. 2F) using Direct Coexistence^84,110^ (DC) simulations (details in the SM). The performance of five residue-resolution coarse-grained models (HPS^111^, HPS-cation-*π*^28^, CALVADOS2^98^, Mpipi^76^, and Mpipi-Recharged^105^) is compared against experimentally reported coexistence lines^65^. The sequences of the proteins investigated are included in Table S2 of the SM. As shown in Figs. 1D-F and 2D-F, Mpipi-Recharged, Mpipi, and CALVADOS2 closely match the experimental trends. While the HPS model offers reasonable predictions for the -9F+3Y, -3R+3K, and -6R+6K variants, the HPS-cation-*π* model significantly underperforms in predicting phase behaviour across all variants (dark yellow data in the figures). Additionally, our results demonstrate a moderate decrease in condensate stability when aromatic residues are mutated to phenylalanine, compared to WT+NLS and allY sequences. This underscores the higher energetic gain of Y–Y interactions over F–F contacts, as shown in our previous atomistic potential of mean force simulations and phase diagrams^76^. Moreover, substituting arginine with lysine notably reduces condensation propensity, consistent with experimental observations^63,65^.

We now calculate the critical temperature and critical density from our phase diagrams from DC simulations by using the law of rectilinear diameters and critical exponents^112^ (Eqs. S11 and S12 in the SM). In Figs. 1G-I and 2G-I, we represent the deviation of the critical temperatures obtained for the studied models with respect to the experimental values (taken from Refs.^63,65^ using the WebPlotDigitizer software) for all the studied variants. Interestingly, this representation clarifies which models accurately capture the critical temperature, showing a notable disagreement with the experimental *in vitro* values for the HPS model for some of the variants except for -9F+3Y, -3R+3K and -6R+6K (red bars) and all variants for the HPS-cation-*π* (yellow bars) model. Indeed, a difference of ∼ 10% in T_*c*_ implies a variation of around 30K in absolute temperature. The Mpipi model (cyan bars) systematically overestimates the critical temperature since electrostatic interactions are partially overlooked compared to *π*-*π* and cation-*π* interactions in such model as discussed in Ref.^105^. Finally, CALVADOS2 (lime green bars) and Mpipi-Recharged models (violet bars) accurately describe the phase diagram of the three studied proteins, presenting a difference in T_*c*_ with respect to the experimental values below 3% in most cases.

Single-molecule properties can be related to the collective phase behaviour of proteins^89,111^. In particular, the *θ* temperature (*T*_*θ*_) serves as a quantitative observable of how the temperature affects the single-protein conformation^89^. It is well-known that the conformational ensemble of a given polymer at *T*_*θ*_, is determined by that of an ideal chain^89,113^. The *θ* temperature, as shown in Ref.^89^, can be related to the condensate critical temperature of a protein solution. We have calculated *T*_*θ*_ for the WT+NLS sequence using all the considered models to compare predictions of single-molecule observables versus condensate properties (see section SVII in the SM for further details on this calculation). Notably, our results confirm that there is a consistent correlation between *T*_*θ*_ and *T*_*c*_ (Fig. 3A) obtained from simulations for each model. Nevertheless, that does not guarantee an accurate description of the phase diagram by the model for this sequence.

**FIG 3.**
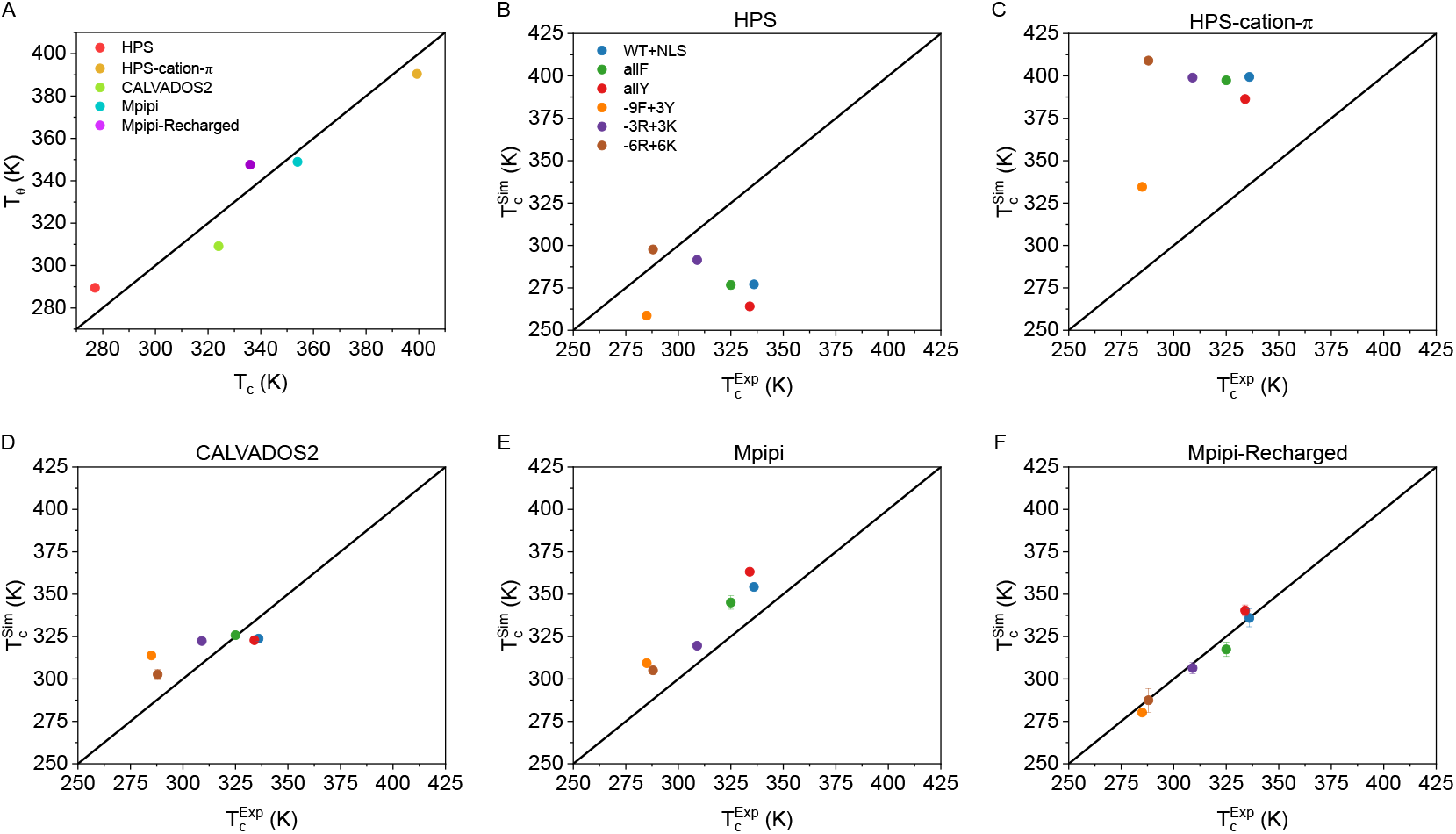
(A) Predicted *θ* temperature (*T*_*θ*_) for the different models *vs*. their critical temperature obtained using the laws of rectilinear diameters and critical exponents for the WT+NLS sequence. Critical temperature predicted by the HPS (B), HPS-cation-*π* (C), CALVADOS2 (D), Mpipi (E), and Mpipi-Recharged (F) models *vs*. the in vitro value of each sequence^63,65^. The black lines indicate a perfect match between experimental and simulated data.

For residue-resolution models, a qualitative description of the phase behaviour of different biomolecules must be applicable to many different protein condensates such as intrinsically disordered proteins^63^, electrostaticdriven biomolecular condensates^69,75^, or multi-domain proteins^5^. In that respect, the models studied in this work sometimes rely on qualitatively capturing the experimental trend so the quantitative description is recovered by applying a scaling factor^28,90^. To explore this, we have compared the critical temperature from our simulations 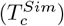 to the experimental critical temperature 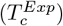 of the WT+NLS variants (Fig. 3B-F) for the different models studied. Interestingly, the HPS and HPS-cation-*π* models do not exhibit a clear trend with respect to the experimentally observed behaviour. In contrast, the CALVADOS2 model provides reasonable predictions of the experimental values, even though the observed correlation is mild. In contrast, the Mpipi and Mpipi-Recharged models accurately capture the experimental trend, with Mpipi-Recharged improving the quantitative predictions of experimental *T*_*c*_.

## III. INTRAMOLECULAR CONTACT MAPS OF HNRNPA1-LCD (WT+NLS) IN DILUTED CONDITIONS

The conformational ensemble of a protein critically determines how it dynamically establishes intra- vs. intermolecular interactions with the surrounding biomolecules. The structural and dynamic behaviour of IDPs can vary widely. IDPs can behave as random coils^114,115^, exhibit disorder-to-order structural transitions^47,48,116^, and even switch among different conformational ensembles^117,118^. Conformational switching refers to the behaviour where intrinsically disordered proteins (IDPs) transition between different conformational ensembles, maintaining partial disorder while fluctuating around an equilibrium structure^119^. This dynamic switching can be regulated by various external factors, such as post-translational modifications like phosphorylation, which has been shown to modulate heterochromatin protein 1 (HP1)^18,70^. Additionally, coarse-grained models suggest that liquid-liquid phase separation can induce a conformational shift in IDPs, driving their transitions from more compact states in dilute conditions to extended conformations in the condensed phase^93,120–^. This behaviour highlights how phase transitions influence the structural dynamics of IDPs during biomolecular condensation.

In this section, we investigate the conformational ensemble of the WT+NLS sequence in the dilute phase (i.e., at the coexistence equilibrium concentration of the solution at T=0.95 T_*c*_) by computing the intramolecular contact maps predicted by the 5 different models (Fig. 4A-E; see section SVI in the SM for further information about this calculation). We find that roughly all models show the standard contact map distribution of an intrinsically disordered polymer, in which most of the interactions are labile, and occur between near neighbouring residues^93^. Indeed, only the Mpipi-Recharged model (Fig. 4F), displays a subtly different map where further neighbours from the diagonal still show significant frequent contacts across the sequence (in green; 75-80th residues contacting the region from the 130-135th residue).

**FIG 4.**
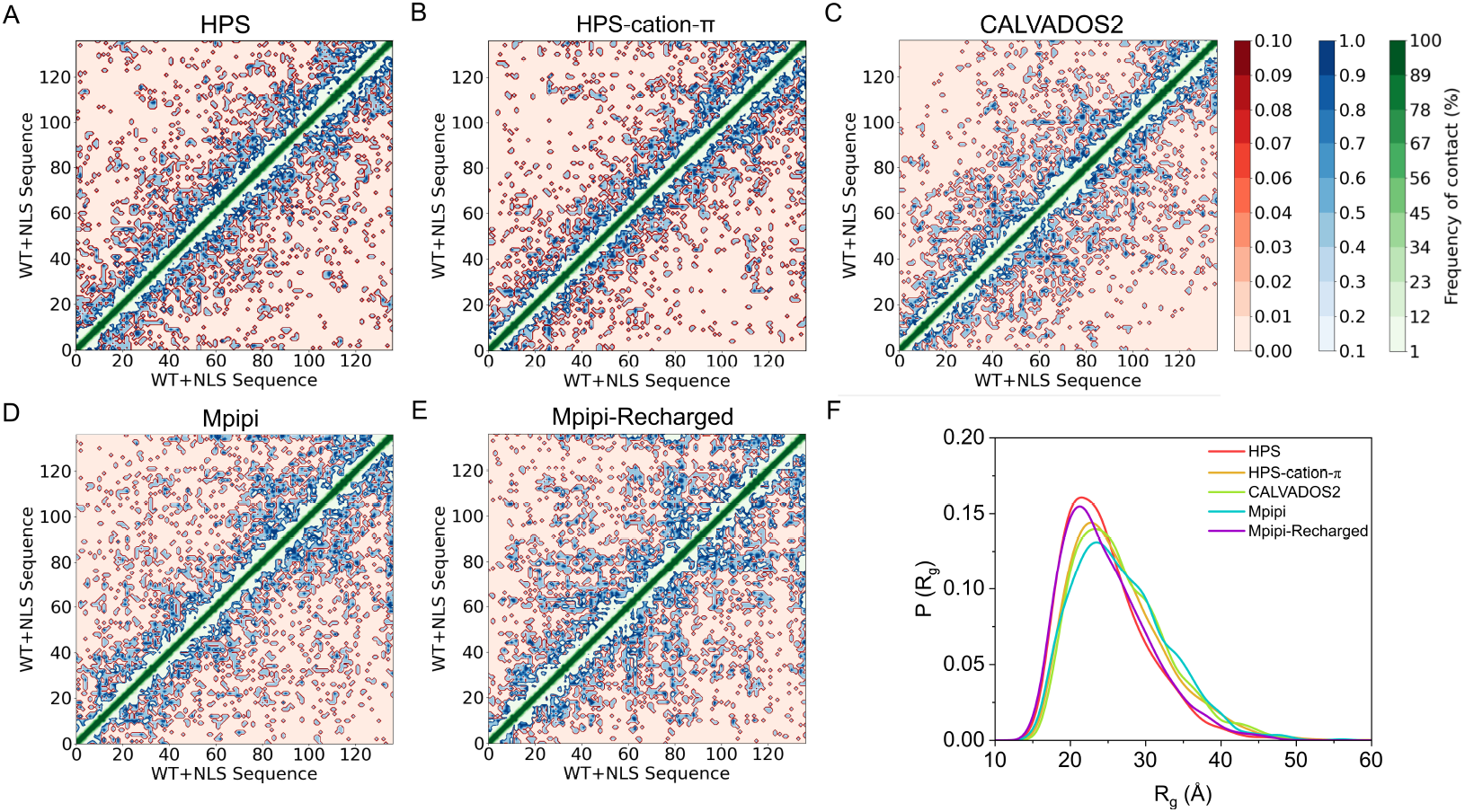
Intramolecular frequency contact maps of the WT+NLS hnRNPA1-LCD variant using simulations of a single protein at T= 0.95 T_*c*_ and at the equilibrium concentration of the diluted phase at such temperature for the HPS (A), HPS-cation-*π* (B), CALVADOS2 (C), Mpipi (D), and Mpipi-Recharged (E) models. The contacts are represented as contact frequency in percentage normalised across the different trajectories generated. Those with frequencies between 100% and 1% are represented in a gradation of green tones, between 1% and 0.1% in blue tones, and those below 0.1% in red tones. (F) Histograms of the radius of gyration of the WT+NLS protein predicted by the different models under the same conditions.

To further investigate the conformational ensemble of the WT+NLS sequence, we now compute the radius of gyration (see further details on this calculation in Section SIV of the SM) of the single-protein in the diluted phase under the same conditions for which the intramolecular contact maps were evaluated. This calculation provides an estimation of the protein compaction, which inherently indicates its propensity to form contacts at intramolecular level^90,121,^. Specifically, we calculate the distribution of the radius of gyration P(*R*_*g*_) of the WT+NLS sequence by the different models at the predicted protein concentration of the dilute phase and at their corresponding values T=0.95 T_*c*_ (Fig. 4F; where T_*c*_ of each model is reported in Table S3). The resulting distributions are consistent with a random coil polymer^90,93^, with an asymmetric curve indicating slight expansion. Notably, despite significant differences in phase diagrams between the models—particularly between the HPS family versus the CALVADOS2 and Mpipi models (Figs. 1 and 2)—the P(*R*_*g*_) distributions remain relatively consistent for all models.

To further investigate this point, we have calculated the radius of gyration (*R*_*g*_) at 300K for all the models and hnRNPA1-LCD sequences (Fig. 5). This enables a quantitative comparison between the model predictions and the experimental values^63^. Fig. 5 shows the average value of the single-molecule protein radius of gyration for the different models (depicted by coloured bars), and the experimental value of each variant is given by the horizontal dashed line. All models, except the HPScation-*π*, provide a reasonable prediction for the value of *R*_*g*_ for all variants, being CALVADOS2 the most accurate model for single-molecule properties. In the case of the HPS-cation-*π* (dark yellow bars in Fig. 5), the radii of gyration are consistently underestimated, and only for the -9F+3Y sequence the model prediction is consistent with the experimental value^63^ (Fig. 5D). Interestingly, the HPS-cation-*π* was parameterized by qualitatively reproducing the experimental trend of the phase behaviour of DDX4 variants^28^, which does not ensure an accurate description of the phase behaviour of other proteins. In fact, other models mainly parameterized for reproducing experimental single-molecule radius of gyration such as the HPS^74^, or other single-molecule properties, such as the CALVADOS2, reproduce well^98^ the experimental values for these variants. Nevertheless, the HPS model does not capture the qualitative trend of the experimental values of *T*_*c*_ for the studied mutants as shown in Figs. 1 and 2. Conversely, both the Mpipi and Mpipi-Recharged models describe reasonably well the radius of gyration of most variants within 0.3-0.4 nm as well as the critical temperature of each sequence. Overall, these results suggest that single-molecule properties, such as the radius of gyration (*R*_*g*_), may be less sensitive to amino acid mutations compared to coexistence lines. While different models can provide reasonable estimates for *R*_*g*_ that align with experimental data (Fig. 5), they may show more pronounced deviations in their predicted phase diagrams (Figs. 1 and 2).

**FIG 5.**
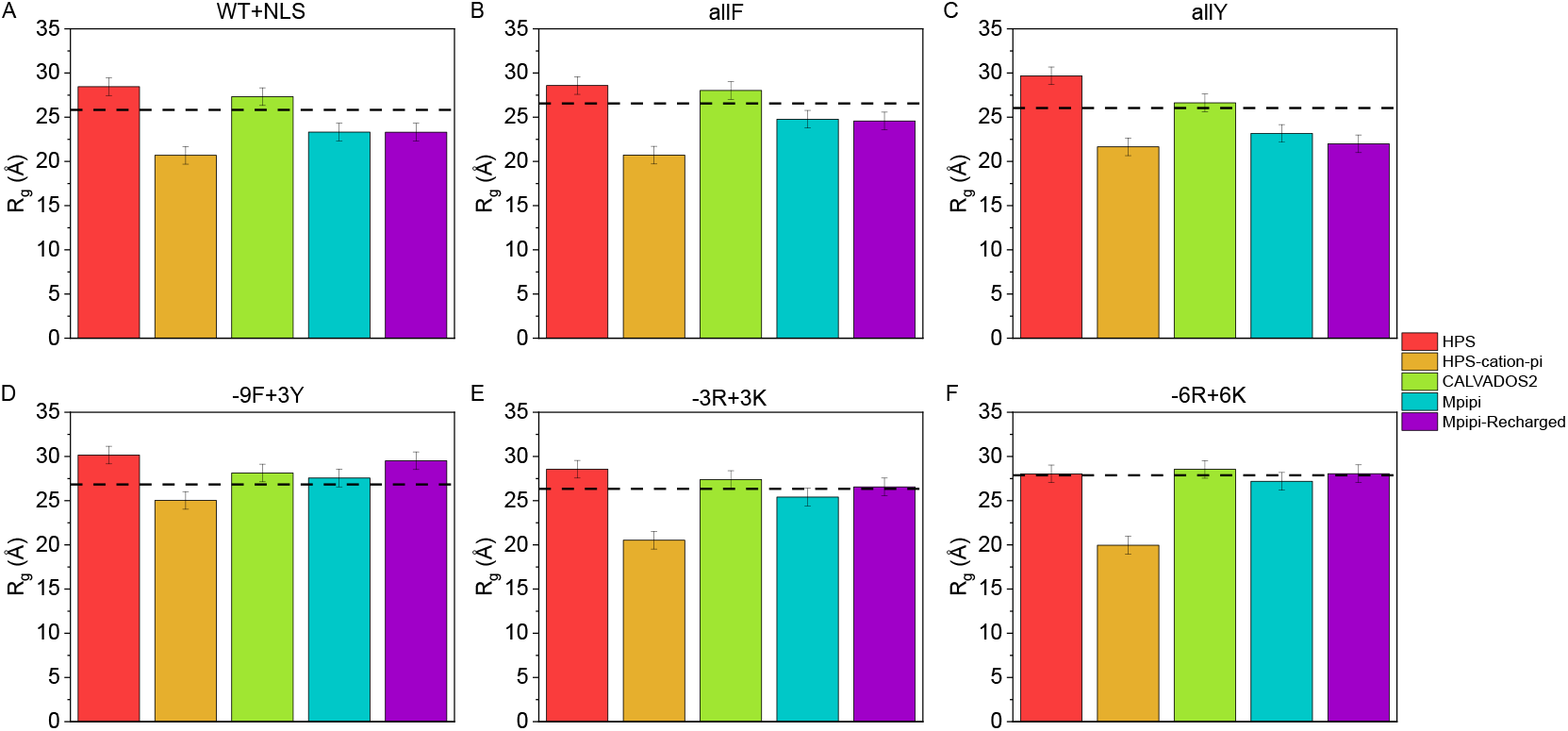
Average single-protein radius of gyration of the different variants WT+NLS (A), allF (B), allY (C), -9F+3Y (D), -3R+3K (E) and -6R+6K (F) predicted by the different models at 300K. The experimentally measured *R*_*g*_^63^ is indicated with an horizontal dashed line.

## IV. KEY INTERMOLECULAR INTERACTIONS FOR WT+NLS ENABLING PHASE SEPARATION

The formation of biomolecular condensates is critically determined by the effective interactions that different biomolecules can establish among themselves^28,90^. Indeed, it is well-known that the presence or absence of aromatic residues in a protein sequence can strongly regulate its phase behaviour^28,62,63,68^. Interestingly, assessing the precise interactions between different proteins inside a condensate is straightforward when using sequence-dependent coarse-grained models^74,76,90,105^, whereas equivalent experimental works (i.e., involving mutagenesis) require the study of the phase behaviour of multiple proteins that differ in few amino acids^62–64,124^. Importantly, the key interactions that sustain biomolecular condensates crucially depend on the specific model and its defined pairwise interactions^74,105^. To better understand the differences between the studied models, we plot their relative interaction strengths of each pairwise interaction (hydrophobic + electrostatic contribution) in Fig. S1 of the SM. As shown, the HPS and CALVADOS2 models present a more homogeneous map for most of the amino acids except for charge-charge interactions, which explains why the variance in the predicted critical temperatures is milder as compared to both Mpipi models (Fig. 3). The HPS-cation-*π* significantly enhances the cation-*π* interactions while keeping the rest of the parameters as the HPS model^28^. Thus, the predicted *T*_*c*_ of the HPS-cation-*π* model are shifted towards higher temperatures compared to the HPS model (Fig. 3B-C). Finally, the Mpipi and Mpipi-Recharged relative interaction strengths present more variance, which explains why their paramaterizations are more sensitive to subtle changes in amino acid mutations (Fig. 3E-F).

We now compute intermolecular frequency contact maps to unravel how different amino acid sequences interact inside the condensates (see section SVI in the SM for further details on this calculation) to overcome the entropic cost of demixing into two coexisting phases^90,96^. We find that different force fields with different relative interaction strength maps (Fig. S1 in the SM) can predict widely different outcomes of how proteins engage with each other to stabilize the condensed phase^59^. In Fig. 6A-E, we compute the intermolecular contact frequency maps between different pairs of amino acids for the WT+NLS sequence using DC simulations at T = 0.95T_*c*_ for the different studied models. Interestingly, using the same distance criterion for all models (section SVI in the SM), the resulting interaction profile heavily depends on each specific parameterization. On the one hand, the HPS model (Fig. 6A) shows a contact map in which a key region from the 85th to the 115th residue presents higher probability of intermolecular contacts with respect to other domains (e.g., the sequence region from the 5th to the 65th residue), that is consistent with the relative interaction map of the model (see Fig. S1 in the SM). On the other hand, the HPS-cation-*π* (Fig. 6B), CALVADOS2 (Fig. 6C), Mpipi (Fig. 6D), and Mpipi-Recharged (Fig. 6E) models show a completely distinct pattern in which specific subdomains display much more prominent contact frequencies than the rest of the sequence. The HPS-cation-*π* predicts the highest contact probability around residues 110-115th and the terminal end beyond residue 130th (Fig. 6B). By inspecting the protein sequence in Fig. 1A and Table S2, we can attribute this result to the accumulation of aromatic and positively charged residues at those particular regions, thus yielding a higher probability of contact between those segments due to the cation-*π* enhancement introduced by this model^28^ (see Fig. S1 in the SM).

**FIG 6.**
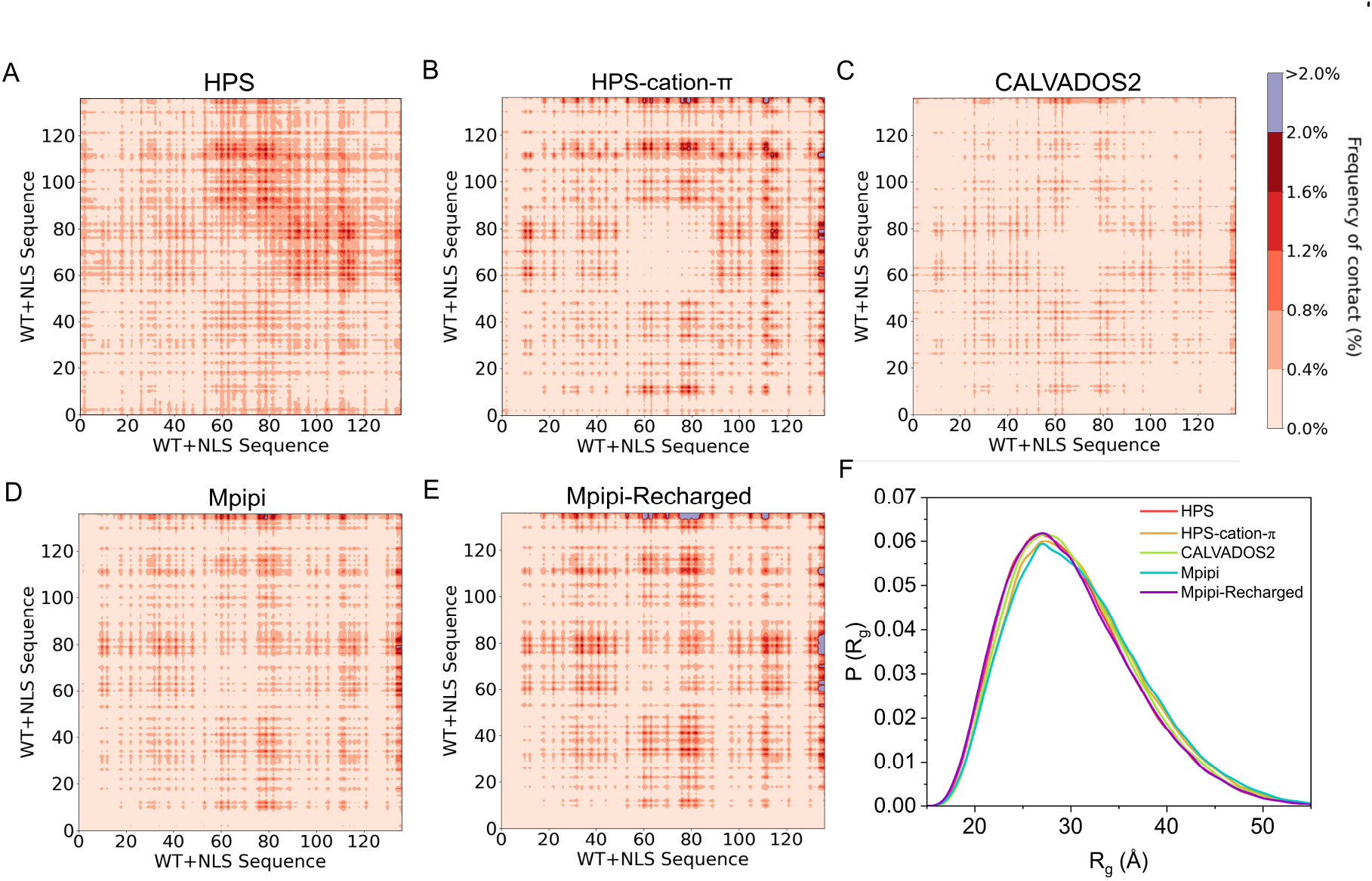
Intermolecular frequency contact maps for the WT+NLS protein from DC simulations at T= 0.95T_*c*_ for the HPS (A), HPS-cation-*π* (B), CALVADOS2 (C), Mpipi (D), and Mpipi-Recharged (E) models. The contacts are represented as contact frequency in percentage. Those with frequencies above 2% have been represented in violet and those below 2% have been represented in a red tone scale. (F) Radius of gyration distributions of the WT+NLS protein predicted by the different models under the same conditions.

The CALVADOS2, Mpipi, and Mpipi-Recharged models depict a similar ‘cross’ motif in their contact maps, which explains their similar prediction of the phase diagram (see Fig. 1D,G). Specifically, the CALVADOS2 model is able to successfully reproduce the phase diagram of the WT+NLS sequence, while its intermolecular frequency contact map exhibits softer variations in contact probabilities across the different sequence subdomains with respect to the Mpipi models (Fig. 6C). Furthermore, for CALVADOS2, the terminal residues (above the 130th) show the highest interaction probability with amino acids located between the 60-80 subdomain, and around the 28-42 region, where negatively charged amino acids are located (Fig. 1A). The Mpipi model, in contrast, predicts larger interaction probability differences between different subdomains along the sequence when compared to the CALVADOS2 model. Notably, the regions with highest probability predicted by the Mpipi correspond to *π*-*π* contacts instead of interactions of electrostatic nature (see Fig. 1A). Finally, the Mpipi-Recharged also suggests *π*-*π* and cation-*π* interactions as the most relevant intermolecular contacts sustaining the WT+NLS LLPS, by predicting a high probability of connectivity between aromatic residues in the terminal part of the sequence (see Fig. 1A). An additional electrostatic contribution for LLPS-stabilising contacts comes from the interactions between the 60-80 subdomain and the terminal region of the sequence (e.g. *>*130th residue) where negatively charged and positively charged residues are located, respectively (see Fig. S1 in the SM). The latter is predicted by the two Mpipi and the CALVADOS2 models.

We now interrogate if the calculation of the radius of gyration within the condensed phase for the different models can provide further information about the intermolecular contacts. Thus, we show the distribution of the radius of gyration (Fig. 6F) calculated under dense phase conditions and at the same temperature as the intermolecular contact maps (*T* ∼ 0.95*T*_*c*_). As shown, the resulting distribution of *R*_*g*_ barely depends on the employed model despite of the dissimilarities exhibited by all of them in their intermolecular contact maps (Fig. 6AE). As expected, and previously found^93,120^, the conformation of WT+NLS in the condensed phase appears significantly more extended than the counterpart in the diluted phase (see Fig. 4F and Fig. 6F). To further analyze such difference, we evaluate the total number of contacts per protein chain within the condensate—both intra-and intermolecular—and compare them with the average intramolecular interactions of the protein in diluted conditions (see section SVI of the SM for further details on this calculation). For intramolecular contacts, we ignore 1–2, 1–3, and 1–4 contacts (i.e., with neighbouring residues below three consecutive adjacent positions). In Fig. 7, the total number of WT+NLS contacts predicted by the different models is represented. We observe that in all models, the number of intramolecular contacts in the dilute phase slightly exceeds those in the condensed phase. This behavior is expected, as single proteins in dilute conditions are known to adopt more compact conformations compared to the condensed phase, as this maximizes their enthalpic gain through intramolecular interactions^90,93,111,120^. This is also consistent with our results for the *R*_*g*_ distribution (Figs. 4F and 6F). However, the total number of contacts within the protein condensate—including both intra- and intermolecular contacts—significantly exceeds the number of contacts in the diluted regime as previously found for multi-domain proteins undergoing phase-separation such as FUS or TDP-43^90^. The five studied models predict a near-quantitatively similar behaviour in that respect, considering the slightly different densities that each model provides at T = 0.95 T_*c*_ (Fig. 1D). Overall from these results we conclude that while all models can provide similar description of the radius of gyration at both single-molecule level and within the condensed phase at a single temperature (Figs. 4F, 5, and 6F), only the CALVADOS2, Mpipi, and Mpipi-Recharged models predict coexistence lines for the hnRNPA1-LCD variants in agreement with experiments (Figs. 1D-I and 2D-I). Reassuringly, despite being developed using entirely different parameterization methods—Mpipi models following a bottom-up approach while CALVADOS2 using a top-down strategy—the three models show consistent agreement in predicting the patterns of intermolecular contacts that sustain condensates.

**FIG 7.**
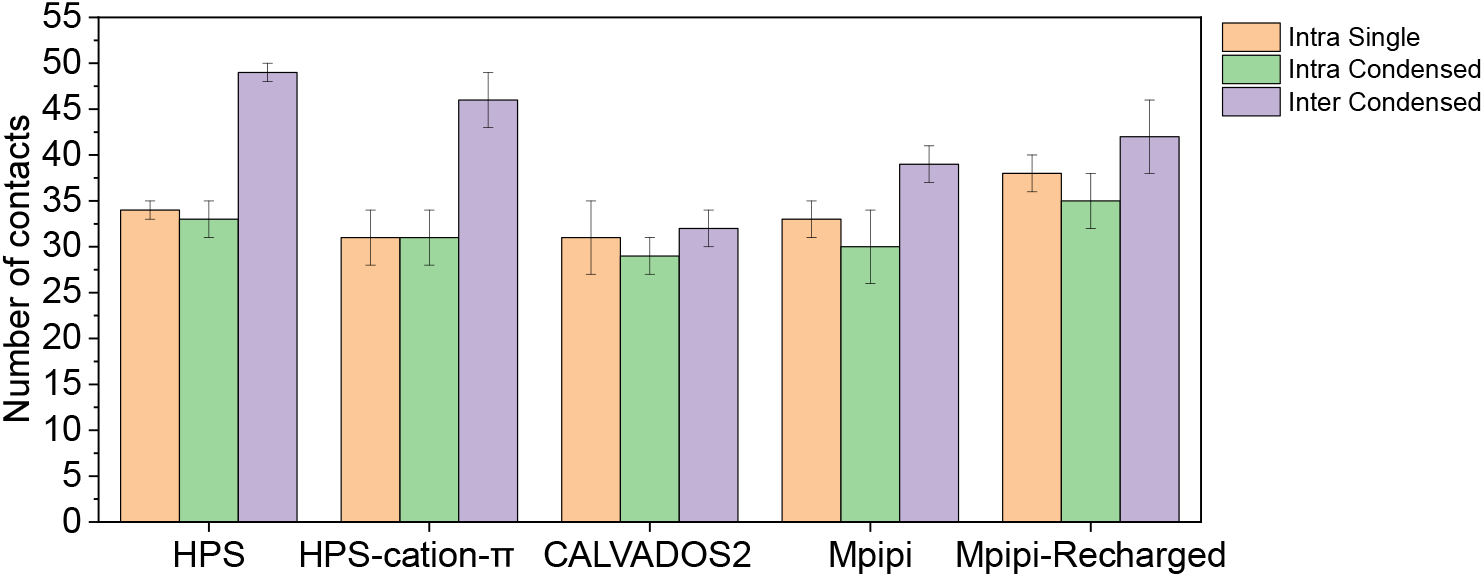
Total number of contacts per protein chain of the WT+NLS sequence at T= 0.95 T_*c*_ predicted by the HPS, HPS-cation-*π*, CALVADOS2, Mpipi, and Mpipi-Recharged models. Orange bars represent intramolecular contacts in simulations with a single protein at diluted conditions; while green and violet bars account for intramolecular and intermolecular contacts, respectively, in simulations of the condensed phase. Intramolecular 1-2, 1-3, and 1-4 consecutive native contacts across the sequence were not considered.

## V. CONCLUSIONS

In this work we analyse and compare the ability of five sequence-dependent coarse-grained models—HPS, HPS-cation-*π*, CALVADOS2, Mpipi, and Mpipi-Recharged—to capture the phase behaviour of six different aromatic-rich intrinsically disordered proteins. We assess the performance of these models by computing the phase diagram of three aromatic variants and three charged variants^63,65^ of the hnRNPA1-LCD sequence in which the identity of a few aromatic and charged residues has been mutated (Fig. 1 and Fig. 2). We find that the mutations explored significantly impact the phase separation propensity of the proteins (Fig. 1D-F and Fig. 2D-F), as experimentally reported^63,65^. However, when comparing to experimental data not all the models are able to capture the experimentally reported trend. While the CALVADOS2, Mpipi, and Mpipi-Recharged models predictions are in near-quantitatively agreement with the experimental phase diagrams for these sequences at physiological salt conditions^65^, the phase diagrams computed with HPS, and especially the HPS-cation-*π* force field, present notable deviations. Moreover, the Mpipi and Mpipi-Recharged models offer the closest predictions to the experimental phase separation propensity of the studied variants (Fig. 3).

We also calculate single-molecule protein properties at diluted conditions including intramolecular contact frequency maps, *θ* temperatures, and the radius of gyration of the WT+NLS sequence to establish a relationship between the protein conformational ensemble at diluted conditions and the intramolecular contact patterning enforcing such dynamical ensemble. In contrast to the predicted phase diagrams, we do not find significant differences neither in the intramolecular contact patterns nor in the radius of gyration distribution predicted by the different models at the same *T/T*_*c*_, which may suggest that a temperature re-scaling could lead to similar predictions by all the models. Remarkably, the predicted value of the radius of gyration at 300K (Fig. 5) agrees within 0.4 nm with the experimental values for most of the variants and models, except for the HPS-cation-*π* model. Furthermore, we find that all models exhibit a higher number of intramolecular contacts per protein in the dilute phase compared to when they are part of a condensate. However, within the condensate, the total number of contacts per protein (intra- and intermolecular combined) is higher than in the dilute phase (Fig. 7).

Regarding the intermolecular frequency contact maps of the WT+NLS condensates at the same *T/T*_*c*_ (Fig. 6), we encounter striking differences in the liquid network connectivity patterns predicted by each model. Whereas the CALVADOS2, Mpipi and Mpipi-Recharged models suggest a similar distribution of intermolecular interactions across the condensate, both HPS and HPS-cation-*π*, suggest an entirely different map of contact frequencies. Our results suggest that single-molecule properties of IDPs can be reliably predicted by residue-resolution coarse-grained models, even when these models differ significantly in their parameters. In contrast, the accurate prediction of phase diagrams is very sensitive to the balance of parameters in the model. Therefore, the accuracy of residue-resolution coarse-grained models for biomolecular phase separation should be evaluated based on collective properties of protein solutions. In particular, given that these models are often paired with the Direct Coexistence method—which enables the calculation of critical temperatures and coexistence densities—the computation of phase diagrams should be considered the gold standard for accuracy testing and benchmarking.

Finally, we note that more sequence-dependent models such as the HPS-Urry^125^ or the FB-HPS^126^ have not been tested in this work due to computational limitations, nevertheless, and as shown in previous work^76^, these models also perform well in describing the phase behaviour of A1-LCD condensates and its variants. In fact, recent models have significantly improved the prediction power in reproducing protein phase behaviour, built upon the influential work of their predecessors^28,74,81^. Overall with this study, we aim to contribute to the continuous improvement and validation of biomolecular coarse-grained models, which can provide unique complementary molecular information that is often hardly accessible through conventional experimental setups.

## VI. ACKNOWLEDGEMENTS

A. F. acknowledges funding from the Ramon y Cajal fellowship (RYC2021-030937-I) and Spanish National Grant (PID2022-136919NA-C33) I. S.-B. acknowledges funding from the Derek Brewer scholarship of Emmanuel College and EPSRC Doctoral Training Programme studentship, number EP/T517847/1. R.C.-G. acknowledges funding from the European Research Council (ERC) under the European Union Horizon 2020 research and innovation programme (grant agreement 803326). A. T. is funded by European Research Council (ERC) under the European Union Horizon 2020 research and innovation programme (grant agreement 803326) and Ramon y Cajal fellowship (RYC2021-030937-I). J. R. E. also acknowledges funding from the Roger Ekins Research Fellowship of Emmanuel College, the Ramon y Cajal fellowship (RYC2021-030937-I) and the Spanish National Agency for Research (PID2022-136919NA-C33). A.R acknowledges funding from PID2023-147156NB-I00 of the Spanish Ministry for Science, Innovation and Universities. This work has been performed using resources provided by the Cambridge Tier-2 system operated by the University of Cambridge Research Computing Service (http://www.hpc.cam.ac.uk) funded by EPSRC Tier-2 capital grant EP/P020259/1.

## Notes

### Competing Interest Statement

The authors have declared no competing interest.

