## Supplementary Material for "Capturing single-molecule properties does not ensure accurate prediction of biomolecular phase diagrams"

Ignacio Sanchez-Burgos

*Maxwell Centre, Cavendish Laboratory, Department of Physics, University of Cambridge, J J Thomson Avenue, Cambridge CB3 0HE, United Kingdom.*

Rosana Collepardo-Guevara

*Yusuf Hamied Department of Chemistry, University of Cambridge, Lensfield Road, Cambridge CB2 1EW, UK and Department of Genetics, University of Cambridge, Cambridge CB2 3EH, UK*

Andrés R. Tejedor†

*Department of Physical-Chemistry Universidad Complutense de Madrid Av. Complutense s/n, Madrid 28040, Spain and Yusuf Hamied Department of Chemistry, University of Cambridge, Lensfield Road, Cambridge CB2 1EW, UK*

(Dated: October 24, 2024)

### SI. MODEL AND METHODS

In these simulations, we reproduce the intrinsically disordered regions (IDRs) as coarse-grained structures with one bead per amino acid. The potential energy of the coarse-grained force field is expressed as follows for each model:

$$E = E_{\text{Bonds}} + E_{\text{Electrostatic}} + E_{\text{Hydrophobic}} + E_{\text{Cation-}\pi} \tag{S1}$$

| Model \ Energy | $E_{\text{Bonds}}$ | $E_{\text{Electrostatic}}$ | $E_{\text{Hydrophobic}}$ | $E_{\text{Cation-}\pi}$ |
|---|---|---|---|---|
| HPS | (S2) | (S3) | (S7) | - |
| HPS-cation-$\pi$ | (S2) | (S3) | (S7) | (S9) |
| CALVADOS2 | (S2) | (S3) | (S7) | - |
| Mpipi | (S2) | (S3) | (S8) | - |
| Mpipi-Recharged | (S2) | (S6) | (S8) | - |

**TABLE S1:** Schematic indices of the formulas describing the interactions subsequently outlined for each model.

In this context, interactions involving $E_{\text{Electrostatic}}$, $E_{\text{Hydrophobic}}$, and $E_{\text{Cation-}\pi}$ forces exclusively involve non-bonded beads. Conversely, $E_{\text{Bonds}}$ accounts for interactions among sequentially bonded beads. In Table S1, schematic indices of the formulas describing the interactions for each model are summarized. The bonding interactions are characterized by a harmonic potential

$$E_{\text{Bonds}} = \sum k(r_i - r_0)^2, \tag{S2}$$

where, $k = 9.6\text{kJ/mol} \cdot \text{Å}^2$ for all the models studied excluding the HPS-cation-$\pi$ whose value is $k = 2.4\text{kJ/mol} \cdot \text{Å}^2$. The equilibrium bond length is $r_0 = 3.81$ Å between bonded amino acid beads in all the cases.

The electrostatic interactions ($E_{\text{Electrostatic}}$) between charged amino acids are governed by a Debye-Hückel potential

$$E_{\text{Electrostatic}} = \sum_i \sum_{j<i} \frac{1}{4\pi\epsilon_r} \frac{q_i q_j}{r} e^{-r\kappa}, \tag{S3}$$

*

†

where $q_i$ and $q_j$ describes the charges of beads $i$ and $j$, $\epsilon_r$= 80 $\epsilon_0$, is the relative dielectric constant of water (being $\epsilon_0$ the dielectric constant). For the CALVADOS2 this constant depends on temperature as shown in Eq. (S9). $r$ is the distance between the $i$th and $j$th beads, and $\kappa = 1$ nm$^{-1}$ is the Debye screening length that mimics the implicit solvent (water and ions) at physiological salt concentration ($c_s \sim$ 150mM of NaCl [1]) for the models HPS, HPS-cation-$\pi$, for the Mpipi model $\kappa = 1.26$ nm$^{-1}$ and for the CALVADOS2 it depends on the salt concentration and temperature as

$$\kappa = \sqrt{8\pi B c_s} \text{ where } B(\epsilon_r) = e^2/4\pi k_B T \epsilon_0 \epsilon_r, \tag{S4}$$

and

$$\epsilon_r(T) = \frac{5321}{T} + 233.76 - 0.9297T + 1.41710^{-3}T^2 - 8.29210^{-7}T^3. \tag{S5}$$

For the Mpipi-Recharged model, the electrostatic interactions are given by a Yukawa potential as

$$E_{\text{Electrostatic}} = \sum_i \sum_{j<i} \frac{A_{ij}}{r} e^{-r\kappa}, \tag{S6}$$

where $A_{ij}$ modulates the interaction between a pair of amino acids and $\kappa$ regulates the salt concentration in an explicit way (Eq. (S4)) and the dielectric constant varies with temperature according to Eq. (S5).

For the HPS, HPS-cation-$\pi$ and CALVADOS2 models, the hydrophobic interactions among amino acid types are designed using a scale of hydrophobicity derived from statistical analysis of amino acid contacts in PDB structures. These interactions are incorporated in a Ashbaugh-Hatch potential [1–4]:

$$E_{\text{Hydrophobic}} = \sum_i \sum_{j<i} \begin{cases} 4\epsilon_{ij} \left[ \left(\frac{\sigma_{ij}}{r}\right)^{12} - \left(\frac{\sigma_{ij}}{r}\right)^{6} \right] + (1-\lambda_{ij})\epsilon_{ij}, & \text{if } r < 2^{1/6}\sigma_{ij} \\ \lambda_{ij} 4\epsilon_{ij} \left[ \left(\frac{\sigma_{ij}}{r}\right)^{12} - \left(\frac{\sigma_{ij}}{r}\right)^{6} \right], & \text{otherwise} \end{cases} \tag{S7}$$

where $\lambda_i$ and $\lambda_j$ are parameters that account for the hydrophobicity of the $i$th and $j$th interacting particles respectively, being $\lambda_{ij} = (\lambda_i + \lambda_j)/2$ following the Lorentz-Berthelot mixing rules [5, 6]. The excluded volume of the residues is given by $\sigma_i$ and $\sigma_j$ for each one, where $\sigma_{ij} = (\sigma_i + \sigma_j)/2$, and $r$ is the distance between the $ij$ particles. $\epsilon_{ij}$ (0.2 kcal/mol) is a fitting parameter to reproduce experimental single-IDR radius of gyration [1].

In the Mpipi and Mpipi-Recharged models, the hydrophobic interactions are given by the Wang-Frenkel potential

$$E_{\text{Hydrophobic}} = \sum_i \sum_{j<i} \epsilon_{ij} \alpha \left( \left[ \frac{\sigma_{ij}}{r} \right]^{2\mu} - 1 \right) \left( \left[ \frac{r_c}{r} \right]^{2\mu} - 1 \right)^{2\nu_{ij}} ,$$
$$\alpha = 2\nu \left( \frac{r_c}{\sigma} \right)^{2\mu} \left[ \frac{1 + 2\nu_{ij}}{2\nu_{ij} \left( \left( \frac{r_c}{\sigma} \right)^{2\mu} - 1 \right)} \right]^{2\nu_{ij}+1} \tag{S8}$$

where the excluded volume of each different residue is given by $\sigma_i$ and $\sigma_j$, where $\sigma_{ij} = (\sigma_i + \sigma_j)/2$, $r$ is the distance between the $ij$ particles, $\epsilon_{ij}$ is the energy interaction term and $r_c$ is the cutoff of the potential between those amino acids, which is always $r_c = 3\sigma_{ij}$, and $\mu$=1 and $\nu_{ij}$ are parameter involved in the shape of the potential.

For the HPS-cation-$\pi$ model another additional component is added in the force field. It includes a reparameterization of the interactions between the positively charged and the aromatic residues given by the Lennard-Jones potential as

$$E_{\text{Cation-}\pi} = 4\epsilon_{ij} \left[ \left( \frac{\sigma_{ij}}{r} \right)^{12} - \left( \frac{\sigma_{ij}}{r} \right)^{6} \right] . \tag{S9}$$

To better visualize the differences in the interactions between the different models we show in Fig. S1 the normalized values of the interaction strength for all the amino acid pairs. The higher the value the more attractive the interaction and vice versa.

### SII. SIMULATION DETAILS

All simulations were carried out using the LAMMPS software [7], in the NVT ensemble, utilizing a Langevin thermostat [8] with a relaxation time of 5 ps. The timestep for the Verlet integration of the equations of motion was set to 10 fs. To enhance computational efficiency, we applied a cutoff of $3\sigma_{ij}$ for the hydrophobic interactions and 3.5 nm for the electrostatic interactions [1]. Each system contained two hundred protein replicas.

### SIII. SEQUENCES OF THE HNRNPA1-LCD AND THEIR VARIANTS

Table S2 presents the sequences of the hnRNPA1 protein and its mutated variants utilized in this work.

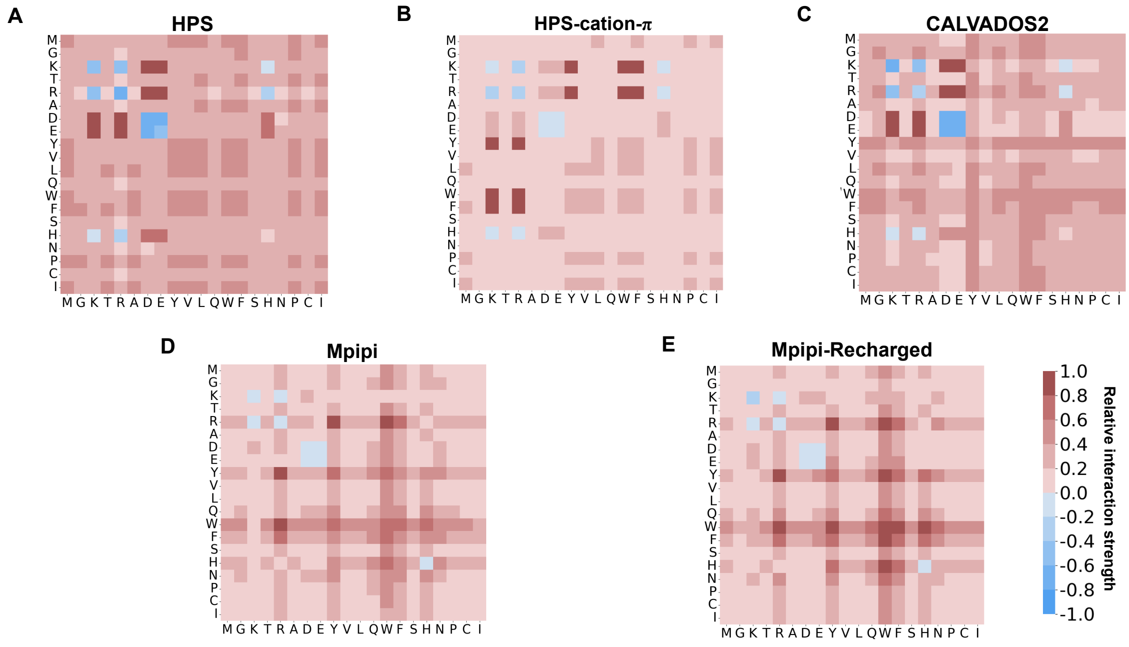


**FIG. S1:** Predicted relative interaction strength values for the HPS (A), HPS-cation-$\pi$ (B), CALVADOS2 (C), Mpipi (D), and Mpipi-Recharged (E) models. The values have been normalized by the highest interaction value of each model.

### SIV. CALCULATING RADIUS OF GYRATION

We analyzed the single-protein conformational ensemble of the different sequences within the diluted phase using the radius of gyration as

$$R_g^2 = \frac{1}{M}\sum_i m_i(r_i - r_{cm})^2. \tag{S10}$$

For this purpose, we compute the distribution of the radius of gyration ($R_g$) of the proteins in the phase. To measure $R_g$, we run NVT simulations with a single protein of the corresponding A1-LCD variant at a temperature equal to $0.95T_c$ of the critical temperature for each model or at 300K for Fig. 5 in the main text. Upon reaching equilibrium, we compute the $R_g$ histograms along the simulation. To calculate the radius of gyration in the condensed phase, we utilized the simulations from direct coexistance simulation at $T = 0.95T_c$ for each model. The result is shown in Fig. 6 of the main text.

| Variant name | Protein sequence | Number and type of change |
|---|---|---|
| **WT+NLS** | GSMASASSSQRGRSGSGNFGGGRGGGFGGNDNFGRGGNFSGRGGFGG<br>SRGGGGYGGSGDGYNGFGNDGSNFGGGGSYNDFGNYNNQSSNFGPMK<br>GGNFGGRSSGPYGGGGQYFAKPRNQGGYGGSSSSSSYGSGRRF | |
| **allY** | GSMASASSSQRGRSGSGNYGGGRGGGYGGNDNYGRGGNYSGRGGYGG<br>SRGGGGYGGSGDGYNGYGNDGSNYGGGGSYNDYGNYNNQSSNYGPMK<br>GGNYGGRSSGGSGGGGQYYAKPRNQGGYGGSSSSSSYGSGRRY | 19Y |
| **allF** | GSMASASSSQRGRSGSGNFGGGRGGGFGGNDNFGRGGNFSGRGGFGG<br>SRGGGGFGGSGDGFNGFGNDGSNFGGGGSFNDFGNFNNQSSNFGPMK<br>GGNFGGRSSGGSGGGGQFFAKPRNQGGFGGSSSSSSFGSGRRF | 19F |
| **-9F+3Y** | GSMASASSSQRGRSGSGNFGGGRGGGYGGNDNGGRGGNYSGRGGFGG<br>SRGGGGYGGSGDGYNGGGNDGSNYGGGGSYNDSGNGNNQSSNFGPMK<br>GGNYGGRSSGGSGGGGQYGAKPRNQGGYGGSSSSSSYGSGRRS | 3F+10Y |
| **-3R+3K** | GSMASASSSQRGKSGSGNFGGGRGGGFGGNDNFGRGGNFSGRGGFGG<br>SKGGGGYGGSGDGYNGFGNDGSNFGGGGSYNDFGNYNNQSSNFGPMK<br>GGNFGGRSSGGSGGGGQYFAKPRNQGGYGGSSSSSSYGSGRKF | 3K |
| **-6R+6K** | GSMASASSSQKGKSGSGNFGGGRGGGFGGNDNFGKGGNFSGRGGFGG<br>SKGGGGYGGSGDGYNGFGNDGSNFGGGGSYNDFGNYNNQSSNFGPMK<br>GGNFGGKSSGGSGGGGQYFAKPRNQGGYGGSSSSSSYGSGRKF | 6K |

**TABLE S2:** The amino acid sequences of the hnRNPA1-LCD variants used in this study are provided. The first column lists the names of the variants, the second column shows their amino acid sequences, and the third column details the number and type of changes compared to the wild-type sequence.

### SV. DIRECT COEXISTENCE TECHNIQUE

With the use of Direct Coexistence (DC) simulations [9–11], we computed the phase diagram for each protein sequence studied [12]. This technique involves simulating two coexisting phases within the same simulation box. In our method, we have placed a high-density protein liquid next to a very low-density counterpart. To accommodate the different densities, we have employed an elongated simulation box, allowing both coexisting phases to form an interface perpendicular to the long axis of the box. Equilibrium has been achieved through NVT simulations, after which we have measured the equilibrium coexisting densities of both phases along the elongated side of the box. This procedure has been repeated at various temperatures until the critical temperature is reached. To reduce finite system-size effects near the critical point, we have calculated the critical temperature ($T_c$) and density ($\rho_c$) using the law of critical exponents and rectilinear diameters [13]

$$(\rho_l - \rho_v)^{\alpha} = s_1 \left(1 - \frac{T}{T_c}\right), \tag{S11}$$

$$\frac{\rho_l + \rho_v}{2} = \rho_c + s_2(T_c - T), \tag{S12}$$

where the notation $\rho_\mathrm{l}$ and $\rho_\mathrm{v}$ represents the densities of the condensed and diluted phases, respectively. In addition, $\mathrm{s}_1$ and $\mathrm{s}_2$ are fitting parameters, while the critical exponent, $\alpha =$ 3.06 from the three-dimensional Ising model [13].

| Variant / Model | WT+NLS | allF | allY | -9F+3Y | -3R+3K | -6R+6K |
|---|---|---|---|---|---|---|
| HPS | 277 K | 277 K | 264 K | 259 K | 291 K | 298 K |
| HPS-cation-$\pi$ | 399 K | 397 K | 386 K | 335 K | 400 K | 409 K |
| CALVADOS2 | 324 K | 326 K | 323 K | 314 K | 322 K | 303 K |
| Mpipi | 354 K | 345 K | 363 K | 309 K | 320 K | 305 K |
| Mpipi-Recharged | 336 K | 317 K | 340 K | 280 K | 306 K | 287 K |
| Experimental | 336 K | 325 K | 334 K | 285 K | 309 K | 288 K |

**TABLE S3:** Critical temperatures of the different variants studied WT+NLS, allF, allY, -9F+3Y, -3R+3K and -6R+6K for the HPS, HPS-cation-$\pi$, CALVADOS2, Mpipi, Mpipi-Recharged models and experimental ones.

### SVI. CALCULATING CONTACT FREQUENCY MAPS AND SUM OF NATIVE CONTACTS

The calculation of intramolecular and intermolecular contact frequency maps per protein chain is derived from single-chain simulations of the protein and DC trajectories for the condensates, respectively (as shown in Fig.4A-E and 6A-E). Contacts have been determined across all systems at temperatures approximately 0.95 times the critical temperature ($T_\mathrm{c}$) of the WT+NLS variant for each respective model. Typically, molecular contacts are determined using a distance criterion, with the assumption that the relative frequency of contact map occurrences (rather than the absolute frequency) remains generally consistent regardless of the chosen cut-off distance, provided it is within a reasonable range. However, to accurately capture the most significant and frequent residue-residue contact pairs that promote LLPS, it is crucial to consider the specific parameterization of each amino acid pair in terms of excluded volume and minimum potential energy interaction distance. Therefore, we have employed an approach, using a sequence-dependent cut-off distance equivalent to $1.2\sigma_{ij}$, where $\sigma_{ij}$ denotes the mean excluded volume of the ith and jth amino acids. Since the potential minimum occurs at approximately $2^{1/6}\sigma_{ij} \approx 1.122\sigma_{ij}$, we set the cut-off distance slightly beyond this point, at $1.2\sigma_{ij}$, to ensure significant binding. By implementing this sequence-dependent cut-off scheme for each amino acid pair interaction, we can effectively filter out adjacent contacts that may coincide with actual interacting amino acids along the sequence, thereby enhancing our ability to accurately identify the amino acids that positively contribute to stabilizing condensates [14].

For the total sum of native contacts in each system, as shown in Fig. 7, we have established a criterion for intramolecular contacts, where we counted only the contacts between amino acids with positions of $i$ and $j \geq i+4$ to avoid counting amino acids that are connected by a bond or a virtual angle in the sequence as a native contact. For intermolecular native contacts, we have considered all existing contacts according to the previously explained criterion.

### SVII. CALCULATION OF $\theta$ TEMPERATURE

To determine the $\theta$ temperature, single-chain simulations were performed across a range of temperatures. For each temperature, as shown in Fig. S2A, we estimated $\nu$, is the Flory scaling exponent, by fitting to

$$R_{ij} = b\,|i - j|^{\nu}, \tag{S13}$$

where $b = 0.55$ nm is the Kuhn length and $i$ and $j$ represent the positions of the amino acids along the sequence. We estimated $T_\theta$ by interpolating the temperature at which $\nu = 0.5$, as shown in Fig. S2B.

[1] G. L. Dignon, W. Zheng, Y. C. Kim, R. B. Best, and J. Mittal, Sequence determinants of protein phase behavior from a coarse-grained model, PLoS computational biology **14**, e1005941 (2018).

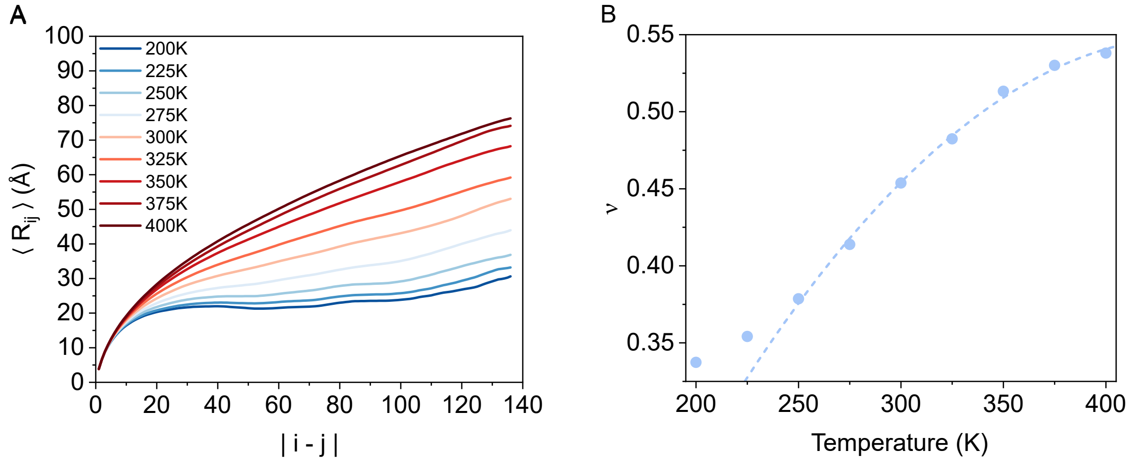


**FIG. S2:** A) Scaling of the average intramolecular distances, $\langle R_{ij} \rangle$ vs. sequence separation $|i-j|$, for different temperatures. B) Calculated $\nu$ against temperature. Dashed line shows the fit to the data to interpolate the temperature at which $\nu = 0.5$


[2] L. H. Kapcha and P. J. Rossky, A simple atomic-level hydrophobicity scale reveals protein interfacial structure, Journal of molecular biology **426**, 484 (2014).

[3] R. M. Regy, G. L. Dignon, W. Zheng, Y. C. Kim, and J. Mittal, Sequence dependent phase separation of protein-polynucleotide mixtures elucidated using molecular simulations, Nucleic acids research **48**, 12593 (2020).

[4] H. S. Ashbaugh and H. W. Hatch, Natively unfolded protein stability as a coil-to-globule transition in charge/hydropathy space, Journal of the American Chemical Society **130**, 9536 (2008).

[5] H. A. Lorentz, On the application of the virial theorem in the kinetic theory of gases, Annals of physics **248**, 127 (1881).

[6] D. Berthelot, Sur le mélange des gaz, Compt. Rendus **126**, 15 (1898).

[7] S. Plimpton, Fast parallel algorithms for short-range molecular dynamics, Journal of computational physics **117**, 1 (1995).

[8] T. Schneider and E. Stoll, Molecular-dynamics study of a three-dimensional one-component model for distortive phase transitions, Physical Review B **17**, 1302 (1978).

[9] A. Ladd and L. Woodcock, Triple-point coexistence properties of the lennard-jones system, Chemical Physics Letters **51**, 155 (1977).

[10] R. García Fernández, J. L. Abascal, and C. Vega, The melting point of ice ih for common

water models calculated from direct coexistence of the solid-liquid interface, The Journal of chemical physics **124** (2006).

[11] J. R. Espinosa, E. Sanz, C. Valeriani, and C. Vega, On fluid-solid direct coexistence simulations: The pseudo-hard sphere model, The Journal of chemical physics **139** (2013).

[12] I. Alshareedah, W. M. Borcherds, S. R. Cohen, A. Singh, A. E. Posey, M. Farag, A. Bremer, G. W. Strout, D. T. Tomares, R. V. Pappu, *et al.*, Sequence-specific interactions determine viscoelasticity and aging dynamics of protein condensates, bioRxiv , 2023 (2023).

[13] J. S. Rowlinson and B. Widom, *Molecular theory of capillarity* (Courier Corporation, 2013).

[14] A. R. Tejedor, A. Garaizar, J. Ramírez, and J. R. Espinosa, ‘rna modulation of transport properties and stability in phase-separated condensates, Biophysical Journal **120**, 5169 (2021).
